# Dendritic cell-intrinsic TLR/MyD88 signaling promotes the clearance of a live-attenuated oral vaccine

**DOI:** 10.1101/2025.06.09.658568

**Authors:** Ying He, Ahmed Mohamed Abdelsalam, Caiting Li, Zhaoyang Liu, Huizhou Fan, Guangming Zhong

## Abstract

A genital tract pathogenicity-attenuated *Chlamydia muridarum* mutant, named intrOv, is being developed into an oral vaccine against *C. trachomatis* infection in the genital tract. Although wild-type *C. muridarum* persists in the mouse large intestine for long periods, intrOv is cleared by the group 3 innate lymphoid cells that produce IFNγ (IFNγ^+^ILC3s). The current study aims to reveal the mechanism by which intrOv induces IFNγ^+^ILC3s for its own clearance from the large intestine. IntrOv-induced IL-23 and IL-23 signaling were required for the clearance of intrOv from the large intestine. The intrOv clearance depended on dendritic cells, as depletion of CD11c+ cells reduced both IL-23 and IFN-γ, resulting in the growth of intrOv. Deletion of MyD88 from dendritic cells, but not phagocytes or epithelial cells, rescued the growth of intrOv, indicating a critical role of MyD88 signaling in dendritic cells. Furthermore, TLR2/4 signaling is also essential for inhibiting intrOv, as the deficiency in TLR2/4 fully rescues the growth of intrOv. Both the TLR and MyD88 signaling must be in the same dendritic cells to inhibit intrOv, as the growth of intrOv in the MyD88- or TLR2/4-deficient mice was blocked by only wild-type bone marrow-derived dendritic cells but not dendritic cells deficient in either MyD88 or TLR2/4. Thus, we have demonstrated a dendritic cell-intrinsic TLR/MyD88/IL-23 pathway for recruiting effectors to clear intrOv from the large intestine. The information may help further improve the efficacy and safety of intrOv and guide the design of future mucosal vaccines.

## Introduction

Despite the urgent need for and the extensive efforts in developing an effective vaccine against Chlamydia in humans [1–14], there is still no licensed vaccine for preventing *Chlamydia trachomatis* [7], the causative agent of human Chlamydia. The efforts in searching for a subunit *C. trachomatis* vaccine, motivated by the failed trachoma vaccine trials [1, 2, 15-17], have produced no licensed vaccines thus far [6, 9, 10, 12, 18]. The success in using nanoparticle-modified *C. trachomatis* organisms to induce protection without exacerbating pathology rekindled interest in developing a whole-cell-based Chlamydia vaccine [11]. The mouse-adapted *C. muridarum*-based pathogenesis studies have led to the identification of genital tract pathogenicity-attenuated *C. muridarum* mutants [19–23], some of which induced protection against subsequent infections with wild-type *C. muridarum* in the genital tract or airway following an oral immunization [24–26]. The mutant designated as an <u>intr</u>acellular <u>O</u>ral vaccine <u>v</u>ector (intrOv) is being developed into an oral vaccine to protect against *C. trachomatis* infection in the genital tract.

While oral inoculation with intrOv induces transmucosal protection in extra-gut mucosal tissues [26], intrOv is cleared from the gastrointestinal (GI) tract [27] by IFNγ [28–31]. Using an intracolon inoculation model, it was found that the IFNγ responsible for the clearance of intrOv was delivered by the IFNγ-producing group 3 innate lymphoid cells or IFNγ^+^ILC3s. However, it remains unclear how intrOv induces IFNγ^+^ILC3s. Revealing the mechanism of intrOv induction of IFNγ^+^ILC3s may provide important information for both further improving the safety and efficacy of intrOv and guiding the design of future mucosal vaccines.

IL-23 signaling was found to be critical in intrOv induction of IFNγ^+^ILC3s, as the clearance of intrOv was blocked in mice deficient in IL-23 receptor (IL-23R) but not IL-1R or IL-18R [32]. However, it remained unknown how intrOv induced IL-23. IL-23 is a heterodimeric cytokine consisting of p19, an IL-23-unique subunit, and p40, a common subunit shared between IL-23 and IL-12 (also called IL-12p40). IL-23 binds to a heterodimeric IL-23 receptor (IL-23R) complex consisting of IL-23R and IL-12 receptor subunit β1 (IL-12R β1). The IL-23p19 subunit first binds to the IL-23R chain, allowing the IL-12p40 to then bind to IL-12Rβ1, both of which are required for the IL-23 signaling that leads to the activation of Janus kinase 2 (JAK2) and tyrosine kinase 2 (TYK2). JAK2/TYK2 may phosphorylate mainly STAT3, but also STAT1, STAT4, and STAT5, to trigger gene expression in cells that express the IL-23R complex [33]. The IL-23R complex is constitutively expressed or induced in myeloid and lymphoid cells. Besides its role in promoting the differentiation of conventional T cells into the Th17 subset, IL-23 signaling can also promote the production of IFNγ by unconventional lymphocytes and innate lymphoid cells (ILCs), including ILC3s [34–36]. IntrOv may induce IL-23 to recruit IFNγ^+^ILC3s for its own clearance from the GI tract. Consistently, IL-23 is frequently detected during chlamydial infection [37–40]. However, the cellular basis of the IL-23 production during chlamydial infection remains unclear. In general, IL-23 is produced mainly by tissue-resident myeloid cells in response to tissue injury or pathogenic insults [41]. The TLR/MyD88 signaling pathway has been shown to induce dendritic cells (DCs) to produce IL-23 [42]. The MyD88 signaling in dendritic cells (DCs) but not phagocytes is also required for inducing an early ILC3s response during intestinal infection with *Citrobacter rodentium* [43]. We hypothesize that intrOv may also activate MyD88 signaling in DCs to produce IL-23.

The current study aims to test whether IL-23 signaling is both necessary and sufficient for intrOv induction of IFNγ^+^ILC3s and, if so, to further determine the cellular basis and signaling pathways of IL-23 production. We have found that intrOv induces IL-23 in the large intestine, and the clearance of intrOv from the large intestine depends on IL-23 signaling. The intrOv induction of IL-23 depended on DCs, as depletion of CD11c+ cells both reduced the levels of IL-23 and IFNγ and increased the yields of intrOv. A cell type-specific knockout in combination with an adoptive transfer approach revealed critical roles of TLR2/4 and MyD88 signaling in the production of IL-23 in DCs and the clearance of intrOv from the large intestine. Furthermore, the TLR2/4 and MyD88 signaling pathways must originate from the same DCs to inhibit the growth of intravascular organisms. Thus, the current study has demonstrated a DC-intrinsic TLR/MyD88/IL-23 pathway for recruiting IFNγ^+^ILC3s to control intracellular bacterial infection. The information may help improve the efficacy and safety of the attenuated vaccine intrOv, and inform the design of future mucosal vaccines.

## Results

### 1. IL-23 signaling is required for the ILC3s-dependent clearance of intrOv from the large intestine

We previously demonstrated that intrOv was cleared from the large intestine by IFNγ^+^ILC3s, and mice deficient in IL-23R were defective in inducing IFNγ^+^ILC3 and failed to clear intrOv from the large intestine [28–30]. In the current study, we first validated the role of IL-23 signaling, as it is known to promote ILC3s to produce IFNγ [44, 45]. As shown in Fig.1, IL-23 was significantly induced in the colon tissue on day 7 after intracolonic inoculation with intrOv, which correlated with a significant reduction in the yield of intrOv (Fig.1A). However, mice deficient in p40, a common subunit shared between IL-23 and IL-12, allowed intrOv to grow in the large intestine (Fig.1B), indicating a critical role of IL-23 &/or IL-12 in restricting intrOv replication. The fact that mice deficient in p35, a subunit unique to IL-12, cleared intrOv from the large intestine as efficiently as the wild-type mice demonstrated that IL-23, but not IL-12, is required for the clearance of intrOv. Furthermore, the clearance of intrOv depended on IL-23R signaling in ILC3s, as IL-23R^-/-^ mice were rescued to clear intrOv by IL-23R-competent RORγt^+^ILC3s (Fig.1C). Thus, the clearance of intrOv depends on both IL-23 and IL-23R.

**Fig. 1.**
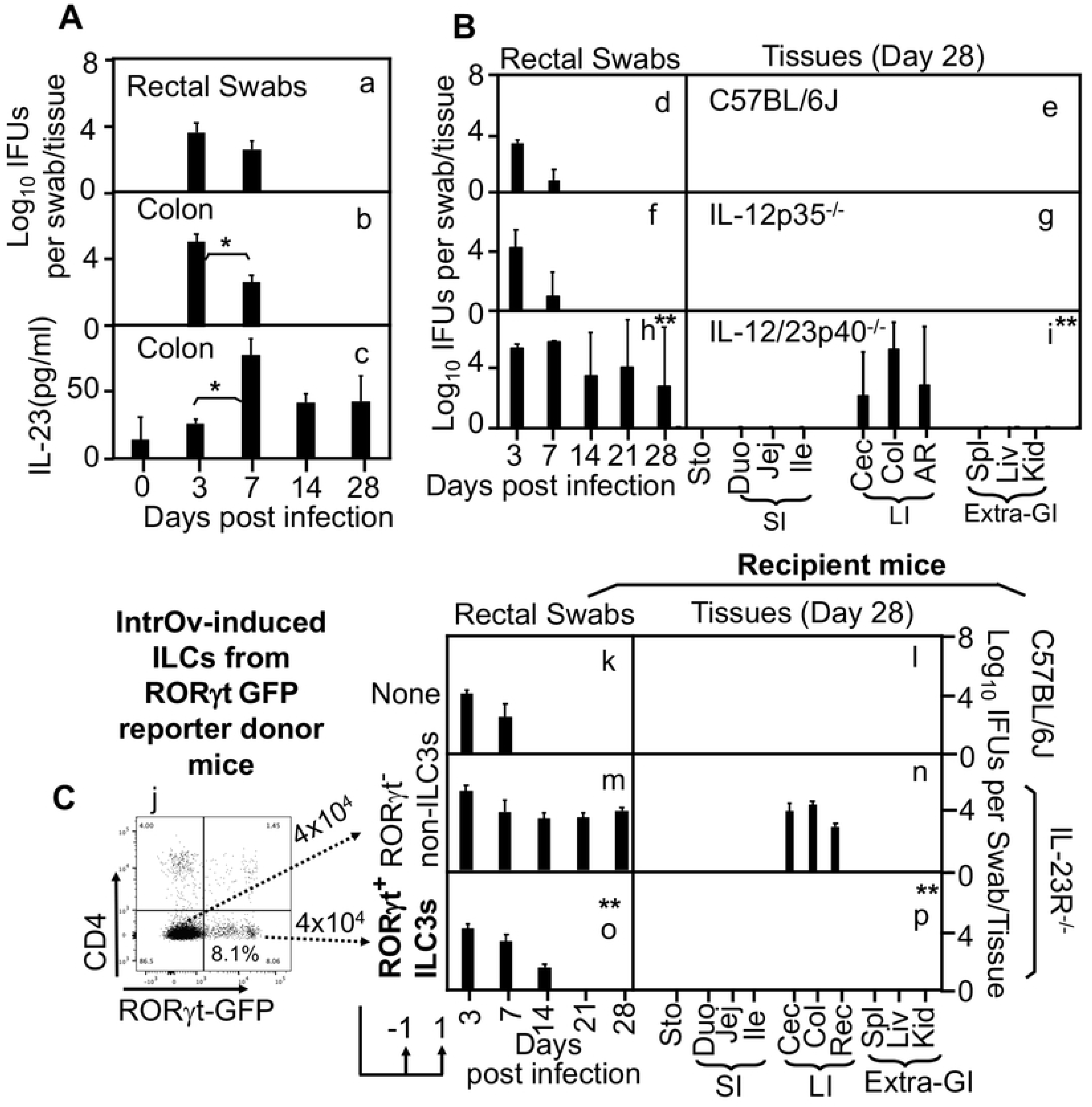
The clearance of intrOv from the large intestine is dependent on IL-23 signaling. A. Mice intracolonically inoculated with 1 x 10^7^ inclusion-forming units (IFUs) of the attenuated oral vaccine strain intrOv were monitored for live intrOv organisms from rectal swabs (panel a, n=5 to 10) or colon tissues (b, n=5) and IL-23 cytokine levels in the colon tissues (c, n=5) on different days post-inoculation (X-axis). Note that IL-23 peaked on day 7, correlating with the reduction in the live intrOv recoveries; B. Mice without (d & e, C57BL/6J, n=5) or with deficiency in IL-12p35 (f & g, IL-12p35^-/-^, n=5) or IL-12p40 (h & I, IL-12p40^-/-^, n=5) were intracolonically inoculated with intrOv and monitored for live intrOv organisms from rectal swabs (d, f & h) or tissues (e, g & i) as described above. Note that IL-12p40^-/-^ but not IL-12p35^-/-^ allowed intrOv persistence in the large intestine; C. Recipient mice without (k & i, C57BL/6J, n=5) or with deficiency in IL-23 receptor (m to p, IL-23R^-/-^, n=5/ea) were intracolonically inoculated with intrOv and monitored for live intrOv from rectal swabs over a time course (k, m & o) or tissues on day 28 (l, n & p) as listed along the X-axis. The IL-23R^-/-^ mice were provided twice (one day before and one day after the intracolon inoculation, respectively) with 4 x 10^4^ RORγt^+^ILC3s (o & p) or RORγt^-^nonILC3s (m & n) sorted from RORγt-GFP reporter mice (j). Note that the RORγt^+^ILC3s but not the nonILC3s donor cells rescued the IL-23R^-/-^ mice to clear intrOv from the large intestine. *p<0.05 (Day 3 vs 7) while **p<0.01 (Area under curves or AUCs), 2-tailed Wilcoxon.

### 2. The induction of IL-23 by intrOv in the large intestine depends on DCs

To determine the cellular basis of IL-23 production during intrOv colonization, we evaluated the role of DCs in the clearance of intrOv, as myeloid cells, particularly activated DCs, are a major producer of IL-23 during microbial infections [41, 42]. When mice expressing the diphtherial toxin receptor (DTR) under the control of the CD11c promoter (CD11c-DTR) were injected with a diphtherial toxin (DT) on even days, the CD11c+ cells were significantly depleted during the 1^st^ week but recovered afterward (Fig. 1S), which is consistent with the known limitation of this approach [46]. Nevertheless, the depletion of DCs during the 1^st^ week was sufficient to significantly reduce the production of IL-23 and IFNγ in the colon on day 7 after intrOv inoculation (Fig. 2A). The reduction in IL-23 and IFNγ correlated with a significant increase in the yield of live intrOv (Fig. 2B). Thus, DCs appear to be essential for both the production of IL-23 and the inhibition of intrOv.

**Fig. 2.**
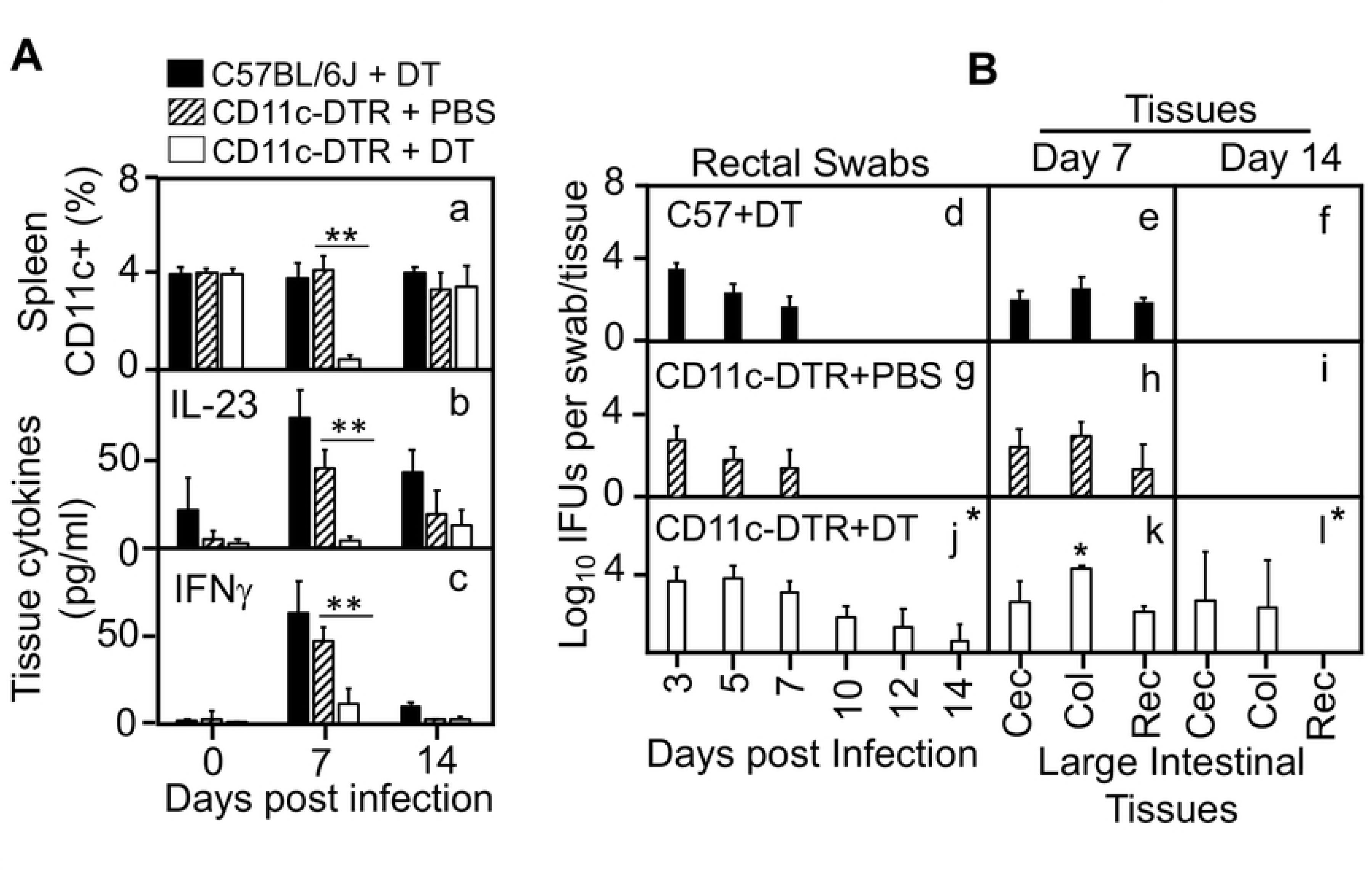
ItrOv induction of IL-23 in the large intestine is dependent on dendritic cells. A. Wild type (solid bar, C57BL/6J, n=5) or transgenic (expressing diphtherial toxin receptor under the control by the mouse CD11c promoter, CD11c-DTR) mice treated with the phosphate-buffered solution (hatched bar, CD11c-DTR + PBS, n=5) or diphtherial toxin (open bar, CDC11c-DTR + DT, n=5) were intracolonically inoculated with intrOv and monitored for CD11c+ cells in the spleen (panel a), IL-23 (b) and IFNγ(c) in the colon tissue on days 7 & 14 after the intracolon infection. Samples from the uninfected mice were used to detect the baselines of IL-23 and IFNγ (day 0). Note that the DT-induced depletion of DCs correlated with the reduction in both IL-23 and IFNγ in the large intestine.B. Wild type treated with DT or transgenic CD11c-DTR mice treated with PBS (n=5) or DT (n=5) were intracolonically inoculated with intrOv and monitored for live intrOv organisms from rectal swabs along the time course (d, g & j) or selected tissues on day 7 (e, h & k) or 14 (f, & l) as described above. Note that DT treatment of CD11c-DTR but not C56BL/6j mice rescued intrOv colonization in the large intestine beyond one week.*p<0.05 while **p<0.01 (between CD11c-DTR mice with or without DT at the single data point or AUCs), 2-tailed Wilcoxon.

### 3. MyD88 signaling in dendritic cells is required for clearing intrOv from the large intestine

To validate the role of DCs and further identify the signaling pathways in the production of IL-23 and the clearance of intrOv, we compared mice deficient in MyD88 from all cells (MyD88^-/-^), CD11c+ cells only (CD11c^Cre^:MyD88^fl/fl^), villin-expressing epithelial cells only (Vill^Cre^:MyD88^fl/fl^) or lymphocytes only (dLCK^Cre^:MyD88^fl/fl^) for their ability to clear intrOv from the large intestine (Fig. 3). The MyD88^-/-^ and CD11c^Cre^:MyD88^fl/fl^ mice allowed intrOv to grow in the large intestine (Fig. 3A). In contrast, the Vill^Cre^:MyD88^fl/fl^ or dLCK^Cre^:MyD88^fl/fl^ mice still cleared intrOv as the wild-type C57BL/6j control mice did. The growth of intrOv correlated with a significant reduction in both IL-23 and IFNγ in the colon of the CD11c^Cre^:MyD88^fl/fl^ mice (Fig. 3B). Thus, DCs and DC-intrinsic MyD88 signaling are required for the production of IL-23 and the clearance of intrOv. Further, the growth of intrOv in the MyD88^-/-^ or CD11c^Cre^:MyD88^fl/fl^ mice was blocked by adoptive transfer with wild-type DCs or IFNγ-deficient DCs (Fig. 4), suggesting although IFNγ is essential for the clearance of intrOv, it is not produced by DCs. Thus, the DC-intrinsic MyD88 signaling but not IFNγ production is necessary and sufficient for DCs to produce IL-23 and promote the clearance of intrOv.

**Fig. 3.**
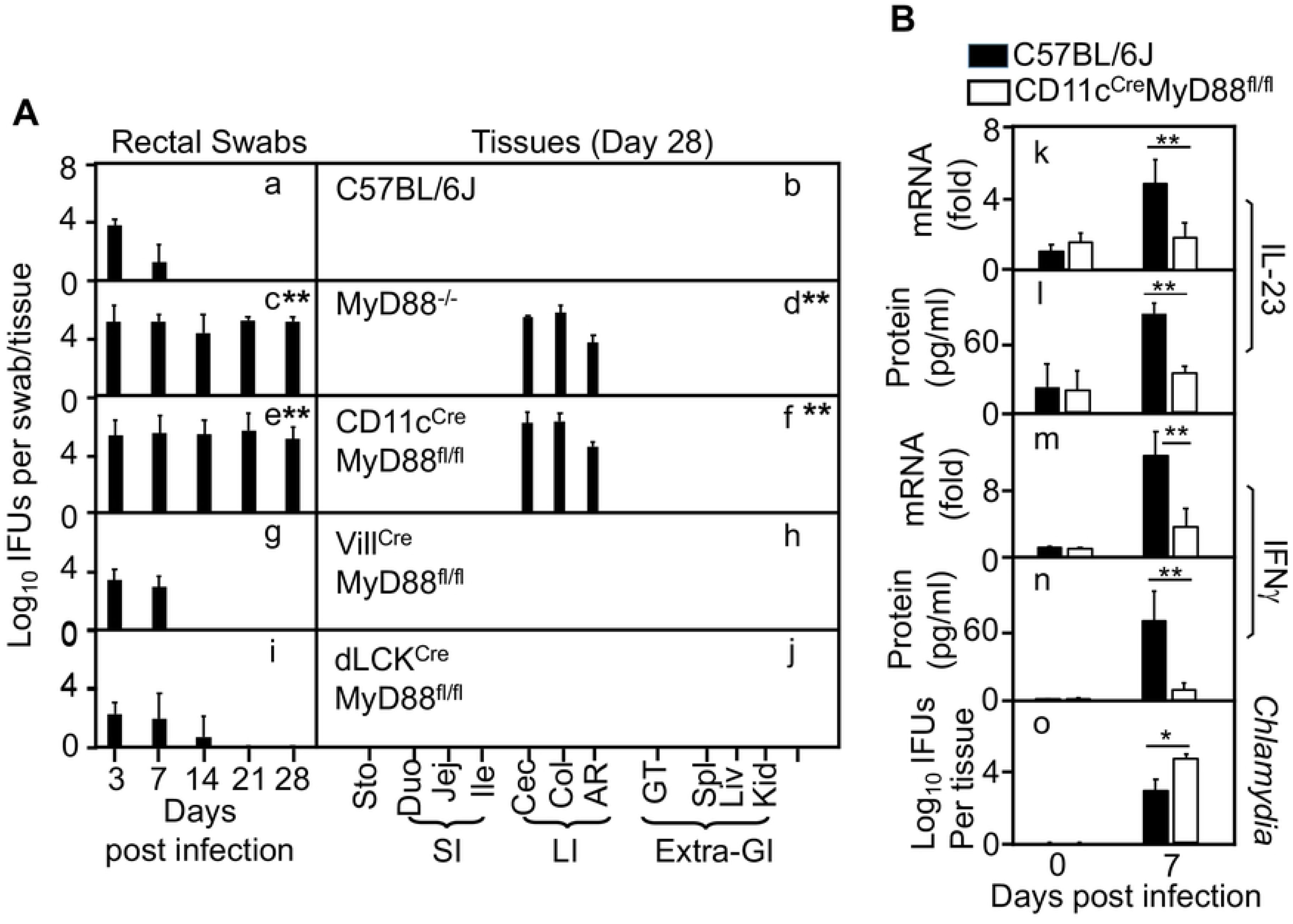
MyD88 signaling in dendritic cells is necessary to clear intrOv colonization from the large intestine. A. Mice without (a & b, C57BL/6J, n=5) or with deficiency in MyD88 from all cells (c & d, MyD88^-/-^, n=5), from CD11c+ cells only (e & f, CD11c^Cre^:MyD88^fl/fl^, n=5), from villin-expressing epithelial cells only (g & h, Vill^Cre^:MyD88^fl/fl^, n=5) or lymphocytes only (i & j, dLCK^Cre^:MyD88^fl/fl^, n=5) were intracolonically inoculated with intrOv and monitored for live intrOv recoveries from rectal swabs (a, c, e, g & i) or tissues (b, d, f, h & j) as described above. Note that only the MyD88^-/-^ and CD11c^Cre^:MyD88^fl/fl^ mice allowed intrOv persistence in the large intestine; B. C57BL/6j control mice (solid bar, n=5) and CD11c^Cre^:MyD88^fl/fl^ mice (open bar, n=5) were intracolonically inoculated with intrOv and monitored for IL-23 mRNA (k) & protein (l), IFNγ mRNA (m) and protein (n) levels as well as the live intrOv titers in the large intestine tissues on day 7 after the intracolon inoculation. The corresponding samples from uninfected mice were used to determine the baselines of the above measurements (day 0). Note that IL-23 and IFNγ were significantly decreased in the KO mice, correlating with the increased the intrOv titers. **p<0.01, (AUVs were compared between C57 and each KO group following ANOVA), 2-tailed Wilcoxon.

**Fig. 4.**
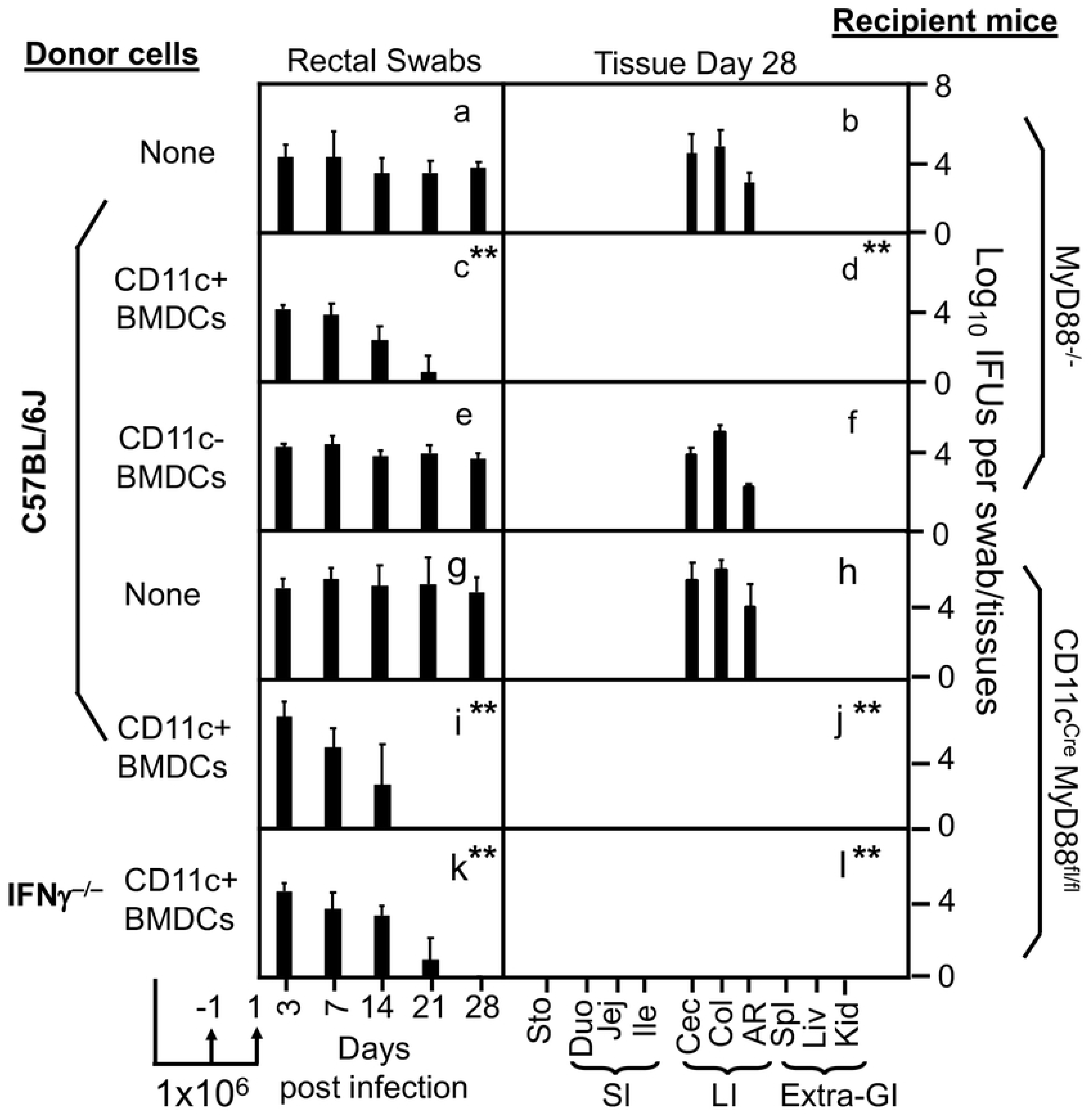
MyD88 signaling in dendritic cells is sufficient for intrOv to induce its clearance from the large intestine. The recipient MyD88^-/-^ (panels a-f) or CD11c^Cre^:MyD88^fl/fl^ (g-l) mice were intracolonically inoculated with intrOv and monitored for live intrOv titers (along Y-axis) from rectal swabs over a time course (a, c, e, g, i & k) or tissues on day 28 (b, d, f, h, j & l) as shown along the X-axis. The MyD88^-/-^ recipients were adoptively transferred twice (one day before and one day after the intracolon inoculation, respectively) each without (panels a & b, None, n=4) or with 1 x 10^6^ CD11c- positive bone marrow-derived cells (c & d, CD11c+BMDCs, n=3) or CD11c-negative bone marrow-derived cells (e & f, CD11c-BMDCs, n=3) harvested from donor C57BL/6j mice. The CD11c^Cre^:MyD88^fl/fl^ recipient mice were similarly without (g & h, None, n=5), or with CD11c+BMDCs from C57BL/6j donor mice (I & j, CD11c+BMDCs, n=5) or IFNγ-deficient donor mice (k & l, IFNγ^-/-^, n=5). Note that CD11c+BMDCs from C57BL/6j or IFNγ^-/-^ donor mice rescued the MyD88-deficient recipients to clear intrOv colonization in the large intestine. **p<0.01, 2-tailed Wilcoxon.

### 4. TLR and MyD88 signaling from the same DCs are required to clear intrOv from the large intestine

Having demonstrated the critical role of DC-intrinsic MyD88 signaling in clearing intrOv, we next identified the upstream receptors. Mice deficient in TLR2 (TLR2^-/-^), TLR4 (TLR4^-/-^), or both (TLR2/4^-/-^), or TLR13 (TLR13^-/-^) were compared for clearing intrOv from the large intestine (Fig. 5). While the TLR2^-/-^ or TLR4^-/-^ mice allowed intrOv to achieve partial growth in the large intestine, the TLR2/4^-/-^ double knockout mice fully rescued the growth of intrOv. The growth of intrOv in the double-knockout mice correlated with a significant reduction in IL-23 and IFNγ, demonstrating a critical role of TLR2/4 signaling in inhibiting the growth of intrOv. Further, the growth of intrOv in TLR2/4^-/-^ or CD11c^Cre^:MyD88^fl/fl^ mice was blocked by adoptive transfer with DCs that are competent in both TLR2/4 and MyD88 but not DCs deficient in either MyD88 or TLR2/4 (Fig. 6). Thus, the TLR2/4 and MyD88 signaling must be in the same DCs to promote the clearance of intrOv from the large intestine.

**Fig. 5.**
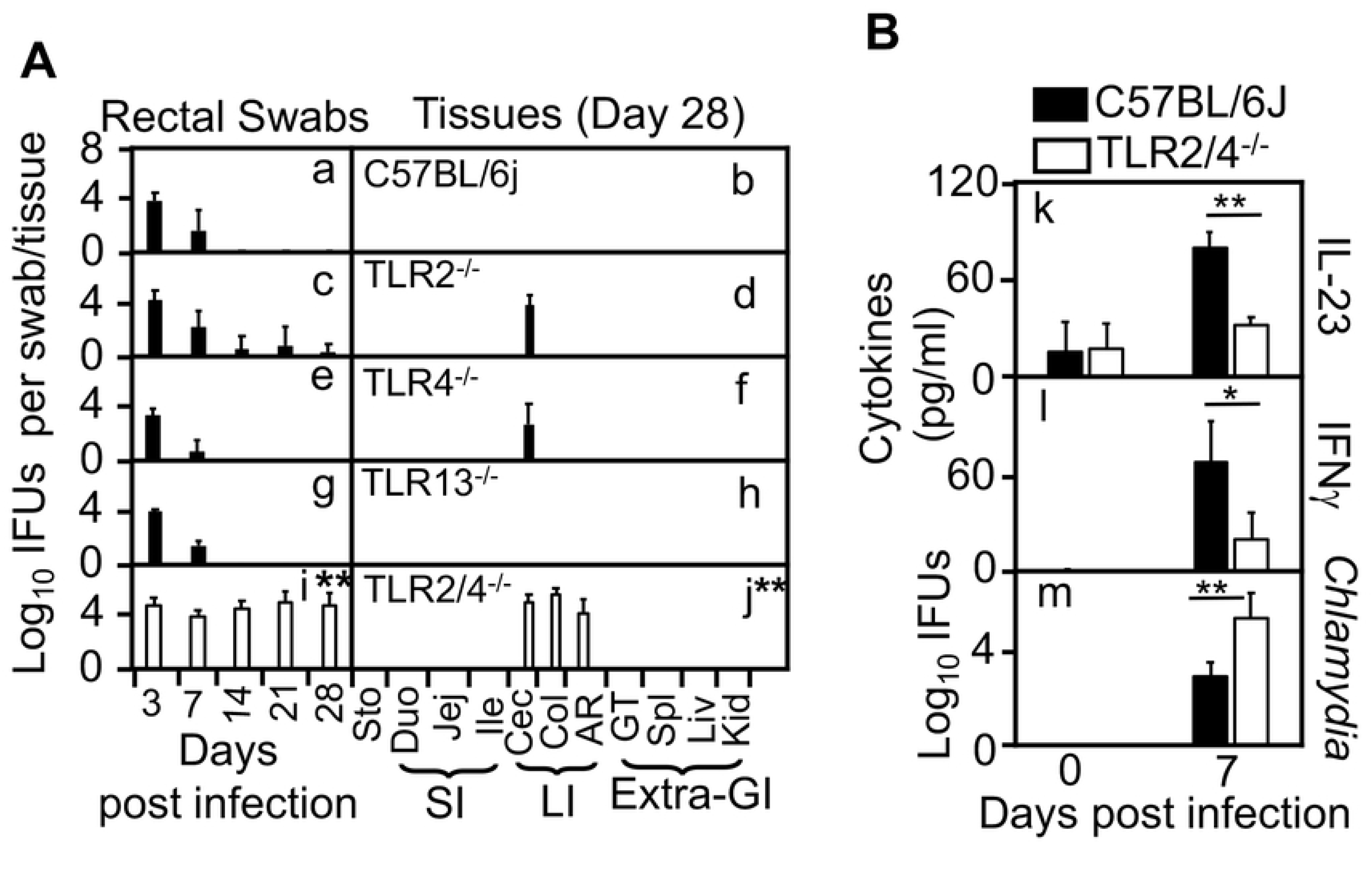
TLR signaling in dendritic cells is necessary to clear intrOv colonization from the large intestine. Mice without (a & b, C57BL/6J, n=5) or with deficiency in TLR2 (c & d, TLR2^-/-^, n=5), TLR4 (e & f, TLR4^-/-^, n=5), TLR13 (g & h, TLR13^-/-^, n=5) or both TLR2 and TLR4 (i & j, TLR2/4^-/-^, n=5) were intracolonically inoculated with intrOv and monitored for live intrOv recoveries from rectal swabs along the infection time course (a, c, e, g & i) or tissues (b, d, f, h & j) as described above. Note that only the double KO TLR2/4-/- mice allowed complete persistence of intrOv in the large intestine, while the TLR2 or 4 single KO mice only permitted partial persistence. B. C57BL/6j (solid bar, n=5) and the TLR2/4^-/-^ double knockout mice (open bar, n=5) were intracolonically inoculated with intrOv and monitored for IL-23 (k) and IFNg (l) protein levels as well as live intrOv recoveries (m) from the large intestine tissues on day 7 after the intracolon inoculation. The corresponding samples from uninfected mice were used to determine the baselines of the above measurements (day 0). Note that IL-23 and IFNγ were significantly decreased in the TLR2/4^-/-^ mice, correlating with the increased the intrOv titers. *p<0.05 while **p<0.01, 2-tailed Wilcoxon.

**Fig. 6.**
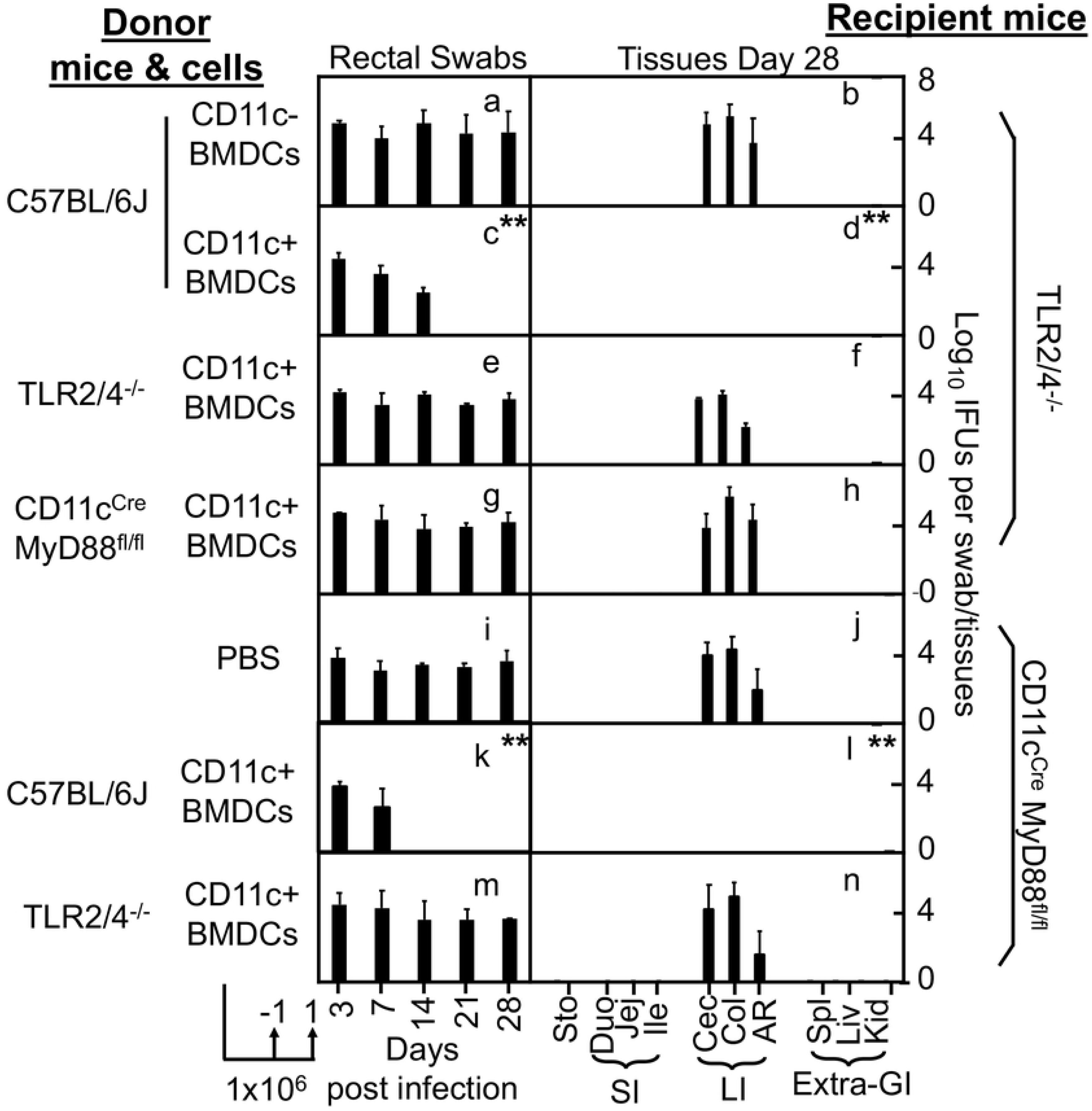
The TLR and MyD88 signaling from the same CD11c+ cells to clear intrOv from the large intestine. The recipient TLR2/4 double knockout mice (panels a-h, TLR2/4^-/-^) or mice deficient in MyD88 from CD11c+ cells only (i to n, CD11c^Cre^:MyD88^fl/fl^) were intracolonically inoculated with intrOv and monitored for live intrOv titers (along Y-axis) from rectal swabs over the infection time course (a, c, e, g, i, k & m) or tissues on day 28 (b, d, f, h, j, l & n) as shown along the X-axis. The TKR2/4^-/-^ recipients were adoptively transferred twice (one day before and one day after the intracolon inoculation, respectively) each with 1 x 10^6^ CD11c-negative bone marrow-derived cells (a & b, CD11c-BMDCs, n=5) or CD11c-positive bone marrow-derived cells (c & d, CD11c+BMDCs, n=5) harvested from C57BL/6j donor mice or CD11c+BMDCs from TLR2/24-/- (e & f, n=5) or CD11c^Cre^:MyD88^fl/fl^ (g & h, n=5). The recipient CD11c^Cre^:MyD88^fl/fl^ mice were adoptively transferred without (I & j, PBS, n=5) or with 1 x 10^6^ CD11c-positive bone marrow-derived cells (CD11c+BMDCs) harvested from C57BL/6j (k & l, n=5) or TLR2/4-/- (m & n, n=5) donor mice. Note that CD11c+BMDCs must be competent in both TLR2/4 signaling and MyD88 signaling to rescue either recipient mice to clear intrOv infection from the large intestine tissues. **p<0.01 (AUCs), 2-tailed Wilcoxon.

## Discussion

IntrOv, an attenuated *C. muridarum* mutant, is currently being developed into an oral vaccine against *C. trachomatis* infection in the genital tract due to both its attenuation in genital tract pathogenicity [21, 22] and capability of inducing transmucosal protection [26]. Because of its susceptibility to IFNγ^+^ILC3s, intrOv is cleared from the GI tract following an oral or intracolonic inoculation [27–31]. The current study has revealed the mechanisms of intrOv induction of IFNγ^+^ILC3s: First, intrOv induced IL-23 and IL-23 signaling was required for clearing intrOv from the large intestine, which is consistent with the previous observations that chlamydial infection induces IL-23 [38–40, 47] and mice deficient in IL-23R failed to inhibit intrOv [32]. Second, the inhibition/clearance of intrOv depended on DCs, as depletion of DCs reduced IL-23 & IFNγ and increased intrOv, demonstrating a critical role of DCs in promoting the clearance of intrOv from the large intestine. Third, deletion of MyD88 from CD11c+ DCs, but not phagocytes or epithelial cells, rescued the growth of intrOv, and adoptive transfer with wild-type bone marrow-derived DCs blocked the growth of intrOv, demonstrating a critical role of DC-intrinsic MyD88 signaling in clearing intrOv. Fourth, mice deficient in TLR2/4 allowed intrOv to grow in the large intestine, and the intrOv growth was inhibited by wild-type DCs, suggesting a critical role of TLR2/4 signaling from DCs. Finally, both the TLR2/4 and MyD88 signaling must be from the same DCs to promote the inhibition of intrOv, as the growth of intrOv in TLR2/4- or MyD88-deficient mice was blocked by adoptive transfer of wild-type DCs but not DCs deficient in TLR2/4 or MyD88. Thus, we have demonstrated a DC-intrinsic TLR/MyD88/IL-23 pathway for recruiting ILC3s to control intracellular bacterial infection in mucosal epithelial cells. The mechanistic information may both help improve the efficacy and safety of the intrOv vaccine, and inform the design of future mucosal vaccines.

ILC3s are highly flexible in differentiating into various stages or subsets with diverse capacities, including the ability to produce IFNγ, becoming IFNγ^+^ILC3s, or ex-ILC3s. IFNγ^+^ILC3s are induced by various enteric infections [48, 49] and can contribute to both protective immunity and pathogenicity [50, 51]. However, the precise mechanism by which each infection induces IFNγ+ILC3s remains unclear. The following receptors, including Ffar2 [52], AhR [53], IL-1R [54], IL-18R [55], and IL-23R [56], may promote ILC3 differentiation in responses to different cues. As an obligate intracellular bacterium variant, intrOv induced IFNγ^+^ILC3s via IL-23R signaling [32]. Consistently, chlamydial infection is known to induce IL-23 [38–40, 47]. The current study has further demonstrated that intrOv can induce IL-23 in the large intestine, and both the intrOv-induced IL-23 and IL-23R signaling on ILC3s are necessary for the clearance of intrOv from the large intestine,

Using a combination of cell type-specific knockout mice with an adoptive transfer approach, we have demonstrated the role of DC-intrinsic TLR/MyD88 signaling in the induction of IL-23 by intrOv and the clearance of intrOv from the large intestine. This finding is consistent with previous observations that IL-23 is mainly produced by myeloid cells [41] and DC-intrinsic TLR/MyD88 signaling is a major pathway for activating DCs to produce IL-23 [42] and induce ILC3s [43]. Although MyD88 signaling has been shown to promote anti-chlamydial immunity [37, 57–59], the current study has provided the 1^st^ comprehensive evidence demonstrating that TLR signaling and MyD88 signaling from the same DCs are both necessary and sufficient for chlamydial induction of IL-23 and anti-chlamydial immunity.

Although intrOv is cleared from the large intestine by the DC-TLR/MyD88-IL-23-IFNγ^+^ILC3s axis, its parental wild type strain *C. muridarum* colonizes the large intestine for long periods [60–63] possibly by evading the immunity mediated by this axis. Microbiota confer colonization resistance against pathogens [64–66]. Maintaining basal levels of type I [67] and type II [65] IFNs by microbiota has been shown to contribute to colony resistance. We hypothesize that the wild-type *C. muridarum*-induction and - evasion of the DC-TLR/MyD88-IL-23-IFNγ^+^ILC3s axis may be a mechanism adapted by *C. muridarum* to maintain a basal level of IFNγ in the gastrointestinal tract. It appears that this basal level of IFNγ is sufficient for clearing intrOv but inadequate for inhibiting the colonization by the wild-type *C. muridarum.* On the one hand, this mechanism may ensure the safety of intrOv as an oral vaccine as it prevents intrOv from continuously shedding live organisms from the gastrointestinal tract. The same mechanism may also contribute to the immunogenicity of intrOv as an oral vaccine. This is because ILC3s represent a subset of RORγt+ antigen-presenting cells [68–71]. The IFNγ^+^ILC3s recruited via the DC-TLR/MyD88-IL-23 axis may present intrOv epitopes to T cells. Efforts are underway to evaluate the contributions of the DC-TLR/MyD88-IL-23 axis-recruited ILC3s as antigen-presenting cells to the transmucosal immunity induced by oral intrOv [26, 32].

## Materials and Methods

### 1. *Chlamydia* organisms

The *Chlamydia muridarum* mutant clone G28.51.1 [21, 22] was used in the current study. This clone is designated intracellular oral vaccine vector or intrOv [72] since it lacks pathogenicity in the genital tract but induces transmucosal protection [26]. IntrOv contains a glutamine (Q) to glutamic acid (E) substitutional mutation at the 117^th^ position of the hypothetical protein TC0237 (TC0237Q117E) and a deletion mutation in the *tc0668 gene,* converting the 216^th^ glycine (G) codon into a premature stop codon (TC0668G216*). The double mutations make intrOv susceptible to IFNγ, and intrOv is cleared from the large intestine by IFNγ-producing group 3 innate lymphoid cells or IFNγ^+^ILC3s [27–30, 32]. The intrOv organisms were grown in HeLa cells (human cervical carcinoma epithelial cells; ATCC# CCL-2), and a density gradient purification protocol was used to purify the elementary bodies (EBs) of intrOv as described previously [73]. The purified EBs were stored in aliquots @-80°C until use.

### 2. Mouse infection and treatment

The mouse experiments were conducted following the recommendations outlined in the Guide for the Care and Use of Laboratory Animals, endorsed by the National Institutes of Health. The protocol was approved by the Committee on the Ethics of Laboratory Animal Experiments of the University of Texas Health Science Center at San Antonio.

The following 6- to 10-week-old male or female mice were used in the current study: C57BL/6J (Jax stock No: 000664, Jackson Laboratories, Inc., Bar Harbor, ME), mice deficient in IL-12p35 (IL-12p35^-/-^; Jax stock No: 002691), IL-12p40^-/-^ (stock No: 002693), IL-23R^-/-^ (035863, *Il23r^tm1Kuch^*/J, IL-23 receptor-eGFP knock-in homozygote), IFNγ^-/-^ (002287), MyD88^-/-^ (009088), TLR2^-/-^ (004650), TLR4^-/-^ (029015). The TLR2/4^-/-^ double knockout mice were bred from the single knockouts. The MyD88^fl/fl^ (008888) mice were crossbred with CD11c^Cre^ (008068) to create CD11c^Cre^-MyD88^fl/fl^ or with Vill^Cre^ (004586) to create Vill^Cre^-MyD88^fl/fl^ or dLCK^Cre^ (012837) to create dLCK^Cre^- MyD88^fl/fl^ mice. The Rorc(γt)-EGFP (007572) heterozygote mice were used as RORγt reporter while CD11c-DTR (004509) mice were used for depleting dendritic cells after injection with diphtherial toxin (DT, cat#D0564, Sigma). The TLR13^-/-^ mice were provided by Dr. Xiaodong Li [74].

All mice were inoculated intracolonically or intragastrically without or with intrOv EBs at a dose of 1 × 10^7^ inclusion forming units (IFUs) per mouse as described previously [26, 30, 75]. Briefly, the intrOv EBs diluted in 50μl of SPG (220 mM sucrose, 12.5 mM phosphate, 4 mM l-glutamic acid, pH 7.5) buffer were delivered to the colon using a straight ball-tipped needle (N-PK 020; Braintree Scientific, Inc., Braintree, MA). After inoculation, mice were monitored for live intrOv organisms in rectal swabs or sacrificed for titrating live organisms and cytokines in organs/tissues, including stomach (STO), various small intestine (SI) segments such as duodenum (Duo), jejunum (Jej), ileum (Ile), various large intestine (LI) segments such as cecum (Cec), colon (Col), anorectum (AR), and extra gastrointestinal (GI) organs such as spleen (Spl), liver (liv), & kidney (Kid). For some experiments, each mouse was intraperitoneally injected with DT at 100ng each on even days for 2 weeks to deplete diphtherial toxin receptor (DTR)-expressing cells.

### 3. Titrating live chlamydial organisms from mouse swabs and tissues

To monitor live chlamydial organisms shedding from the GI tract, rectal swabs were collected in 0.5 ml of SPG buffer and vortexed with glass beads to release infectious EBs. To titrate live organisms from mouse tissues/organs, the corresponding tissues/organs indicated in individual experiments were collected (on designated days specified in the corresponding experiments) into 2 ml of SPG buffer, followed by homogenization and brief sonication. Live intrOv organisms released in the supernatants were titrated in duplicate on HeLa cell monolayers grown in 96-well plates. 50μl of each sample without (neat) or with serial dilutions was used to inoculate monolayer HeLa cells. After incubation overnight, the infected HeLa cells were processed for immunofluorescence labeling of chlamydial organisms as described below. The entire culture well was counted for chlamydial inclusions when the inclusion density was low, allowing the detection of a single IFU per 50μl sample. The total number of IFUs per swab or tissue was converted into log_10_ for calculating the group mean and standard deviation. Typically, the detection limits are 10 IFUs per swab and 40 IFUs per tissue sample.

### 4. Immunofluorescence assay

The immunofluorescence assay for visualizing and counting chlamydial inclusions in the Chlamydia-infected HeLa culture was described previously[76]. Briefly, infected HeLa cells grown on 96-well plates were fixed with paraformaldehyde (Sigma, St. Louis, MO 63178) and permeabilized with Triton X-100 (Sigma). The monolayers were labeled with a rabbit anti-chlamydial antibody (raised by immunization with *C. muridarum* EBs) and a goat anti-rabbit IgG conjugated with Cy2 (green, Jackson ImmunoResearch Laboratories, Inc) to visualize chlamydial inclusions, while a Hoechst dye (blue; Sigma) was used for labeling nuclear DNA. The labeled cell samples were viewed under an Olympus IX-80 fluorescence microscope equipped with multiple filter sets (Olympus, Melville, NY).

### 5. Quantitative reverse transcriptase polymerase chain reaction (qRT-PCR) for detecting mouse *ifnγ* and *IL23p19* expression

For some experiments, mouse tissues were used to measure IFNγ and IL-23 mRNAs and proteins (see ELISA below) in addition to quantifying live intrOv organisms (see above). After sacrificing a mouse, the colon tissue was excised, opened longitudinally, and rinsed off fecal matter. The cleaned tissue was quickly minced, and ¼ of the minced tissue from each sample was aliquoted for RNA extraction, while the remaining tissue pieces were used to make tissue homogenates in SPG as described above. To extract total RNA, the large intestine tissue pieces were further minced before adding to a Lysing Matrix D tube with beads (Cat#: 6913050, MP Biomedicals, Santa Ana, CA). After a quick mix, the sample tube was immediately frozen in liquid nitrogen. Total RNAs were isolated using TRIzol reagent (Cat#: 93289, Sigma) according to the manufacturer’s instructions. 1 ml TRIzol was added to each frozen tube, followed by homogenizing for 45 seconds twice (Mini-Beadbeater, Biospectra, Stroudsburg, PA). After extracting with 200 μL chloroform per tube, the top aqueous phase (about 300-400μl) was carefully collected into a new RNase-free 1.5 mL tube, followed by adding 550μl isopropanol to precipitate nucleic acids. The final sample was resuspended in 20μl DEPC water. First-strand cDNA was synthesized from 200 ng of total RNA in a 20μl reaction using the iScript™ cDNA Synthesis Kit (Cat#: 1708891, Bio-Rad). A SYBR green real-time PCR kit (Cat# 001752A, Bio-Rad) and a CFX96 Real-time Detection System (Bio-Rad) were used to run the qPCR reaction. The 2^-ΔΔ*CT*^ method was used for data analysis [77]. The target genes were mouse *ifnγ* & *il23p19*, respectively, while the reference gene is *gapdh* from the same sample. The following primers were synthesized by Integrated DNA Technologies, Inc. (San Diego, CA) and used in the current study: IFNγ forward primer (5′-TCAAGTGGCATAGATGTGGAAGAA-3’) and reverse primer (5′-TGGCTCTGCAGGATTTTCATG-3’), IL-23p19 forward primer (5’-CCAGCAGCTCTCTCGGAATC-3’) and reverse primer (5’-TCATATGTCCCGCTGGTGC-3’), and GAPDH forward primer (5’-ACCACAGTCCATGCCATCAC-3’) and reverse primer (5’-TCCACCACCCTGTTGCTGTA-3’).

### 6. Measurement of cytokines in mouse tissues using ELISA

The ELISA kits for measuring mouse IFNγ and IL-23p19, respectively, from mouse tissues were purchased from Thermo Fisher Scientific (Cat# 88-7314-88 for IFNγ and Cat# 88-7230-88 for IL-23p19, Waltham, MA). 100 μl of each colon tissue homogenized in SPG buffer as described above was mixed with a protease inhibitor cocktail (cat#78430, 100x stock, Thermo Fisher Scientific) at a final concentration of 2X. The neat homogenates were 2-fold serially diluted with PBS containing 2X inhibitor cocktail, and the serially diluted samples were applied to 96-well plates precoated with a capture antibody. IFNγ or IL-23p19 binding was detected with a detection antibody plus detection reagent. The absorbance at 450 nm was detected with a Synergy H4 microplate reader (BioTek, Winooski, VT), and the results were expressed as pg/ml based on the standard curve obtained from the same plate.

### 7. Adoptive transfer

Innate lymphoid cells (ILC3s) prepared from RORγt-GFP reporter mice or bone marrow-derived dendritic cells (BMDCs) from wild-type or mice deficient in MyD88, TLR2/4, or IFNγ, respectively, as donor cells for adoptive transfer experiments. The transfer was carried out via retro-orbital injection twice, each with a designated number of donor cells, as indicated in individual experiments. The 1^st^ transfer was done on the day before, and the 2^nd^ on the day after the intracolonic inoculation of the recipient mice with intrOv. Following the intracolonic inoculation, live intrOv organisms were monitored in both rectal swabs and tissues.

The donor ILCs were prepared as described previously [30, 78]. Briefly, each donor RORγt-EGFP heterozygote mouse was orally inoculated with 1 x 10^7^ IFUs of intrOv for 7 days before the intestinal tissue was collected for lymphoid cell isolation. After the intraepithelial lymphoid cells were first removed, the remaining tissue pieces were used for isolating lamina propria lymphoid cells by mincing the tissues into 1-2 mm pieces. The minced tissue mixture was digested with both DNase I and Liberase followed by shaking to release lymphoid cells into the supernatants. The process can be repeated several times. After filtering and washing, the lymphoid cell-containing solutions were collected for Percoll purification. The lymphoid cell-containing interphase was harvested as lamina propria lymphoid cells or LPLs for flow cytometry sorting after excluding dead and lineage-positive cells (gating for live lineage-negative or lin^-^ cells). GFP was used to sort for RORγt-expressing cells. The CD4-negative but GFP-positive subset was used as donor ILC3s, while CD4-negative and GFP-negative subset from the same mouse was used as donor non-ILC3s.

The donor bone marrow-derived dendritic cells were prepared from naïve C57 or mice deficient in different genes, as indicated in individual experiments. A standard protocol as described previously [79–81] was followed. Briefly, after the removal of red blood cells, bone marrow cells collected from each donor mouse were cultured for 3 days in one 6-well plate with 3 ml per well and cell density of ∼1 x 10^6^ per ml in RPMI 1640 medium (cat# R8758, Sigma) supplemented with 10% Fetal Bovine Serum (FBS, cat#900-108, Gemini Bio, West Sacramento, CA), 15ng/ml of GM-CSF (cat#50-170-433, Thermo Fisher Scientific), and 10ng/ml of IL-4 (cat#214-14, Thermo Fisher Scientific). On the 3^rd^ day, the floating cells were gently removed, and the partially attached cells were transferred to a new 6-well plate. The cells in the new plate were cultured for 3 additional days in a freshly prepared medium as described above. The 6-day-old cells were used to sort CD11c+ dendritic cells that were collected in PBS as donor cells for adoptive transfer.

### 8. Flow cytometry analysis and sorting

The following antibodies were used for labeling surface markers: rat anti-mouse CD16/32 (block the nonspecific binding to Fc receptors, clone:2.4G2, cat#:BE0307, Bio X Cell, West Lebanon, NH), rat anti-mouse Lineage cocktail (conjugated with Pacific blue, cat#133310, Biolegend, Inc, San Diego, CA) consisting of rat anti-mouse CD3 (clone: 17A2), rat anti-mouse Ly-6G/Ly-6C (clone: RB6-8C5), rat anti-mouse CD11b (clone: M1/70), rat anti-mouse B220 (clone: RA3-6B2) and rat anti-mouse Ter-119 (clone: Ter-119), rat anti-mouse CD45.2 (conjugated with APC, clone#104, cat#17-0454-82, Thermo Fisher Scientific, Inc, Waltham, MA) and rat anti-mouse CD90.2 (conjugated with FITC, clone#30-H12, cat# 105305, Biolegend, Inc). Dead cells were excluded by staining with Viability-Dye (conjugated with eFluor 506, cat#65-0866-14, Thermo Fisher Scientific). The anti-CD4 antibody (clone#RM4-5, cat#BDB553051, Thermo Fisher Scientific) was used to identify conventional CD4+ T cells from the intestinal lamina propria lymphoid prep, while the anti-CD11c antibody (clone#HL3, cat#561241, Thermo Fisher Scientific) was used to identify dendritic cells from BMDC prep. The antibody staining was permitted for 30min at 4°C.

FACS Aria II instrument (BD Biosciences) was used to sort the desired cell population as donor cells for adoptive transfer experiments. After excluding dead and lineage positive cells (CD3+Ly6C+CD11b+B220+Ter119+), GFP+ cells sorted from RORγt-GFP reporter mice were used as donor ILC3s, while GFP-cells were used as non-ILC3 controls. After excluding dead cells, CD11c+ BMDCs were used as donor dendritic cells.

### 9. Statistics

The numbers of live organisms in IFUs at individual data points or over a time course were compared using the Wilcoxon rank-sum test. Area-under-the-curve or AUC was used for comparing the time course or clusters of tissue sample data. When multiple groups were included in a given experiment, ANOVA was first used to determine whether there was an overall significant difference among all groups. Only when p<0.05 (ANOVA), were the differences between every two groups further analyzed using Wilcoxon.

